# Fission stories: Using PomBase to understand *Schizosaccharomyces pombe* biology

**DOI:** 10.1101/2021.09.07.459264

**Authors:** Midori A. Harris, Kim M. Rutherford, Jacqueline Hayles, Antonia Lock, Jürg Bähler, Stephen G. Oliver, Juan Mata, Valerie Wood

## Abstract

PomBase (www.pombase.org), the model organism database (MOD) for the fission yeast *Schizosaccharomyces pombe*, supports research within and beyond the *S. pombe* community by integrating and presenting genetic, molecular, and cell biological knowledge into intuitive displays and comprehensive data collections. With new content, novel query capabilities, and biologist-friendly data summaries and visualisation, PomBase also drives innovation in the MOD community.

## Introduction

Over the past decade, PomBase (www.pombase.org), the authoritative model organism database (MOD) for the fission yeast *Schizosaccharomyces pombe*, has supported the fission yeast research community by integrating and presenting all types of genetic, molecular, cell biological, and systems-level knowledge relevant to *S. pombe*. In addition to about 200 laboratories dedicated to fission yeast research, PomBase serves a growing number of users who focus on other organisms but rely on data generated in *S. pombe* for inferences about orthologous genes and conserved eukaryotic cell biology. As fission yeast research has grown in complexity and breadth of relevance, PomBase has kept pace, supporting emerging experimental and data-handling techniques, enabling researchers to ask novel questions, and driving innovation in the MOD community.

PomBase’s primary aims remain to standardise, integrate, and display fission yeast research, to disseminate datasets and new knowledge to the wider scientific community, and to highlight the added value these efforts bring.

The core of PomBase consists of a growing body of comprehensive, reliable knowledge derived by manual curation of the fission yeast literature. Manual curation covers a wide range of data types, including molecular functions, biological processes, cellular locations, macromolecular complexes, phenotypes, alleles and genotypes, protein modifications, physical interactions, genetic interactions, DNA and protein sequence features, and orthologs in human and budding yeast (*Saccharomyces cerevisiae*). Annotations generated by computational methods supplement manually curated data for function, process, and location data represented using the Gene Ontology (GO; The Gene Ontology Consortium 2000, 2021). Table 1 summarises annotation types and totals.

**Table 1.** Summary of annotations in PomBase as of 1 September 2021.

| Annotation type | Annotation count |
| --- | --- |
| Gene Ontology | 39212 |
| Phenotype | 98966 |
| Protein modification | 48603 |
| Gene expression | 33754 |
| Genetic interactions | 3819 |
| Physical interactions | 2384 |
| Other | 57159 |
| **Total** | **283897** |

Totals exclude interaction annotations imported from BioGRID (Oughtred et al., 2021) and approximately 3200 non-redundant GO annotations based on computational methods. Additional details, and up-to-date totals, are available at https://curation.pombase.org/pombe/stats/annotation.

Literature curation makes extensive use of ontologies, including GO, the Sequence Ontology (SO; Eilbeck et al. 2005), the Protein Ontology (PRO; Natale et al. 2017), the Mondo Disease Ontology (developed by the Monarch Initiative; Mungall et al. 2017; Shefchek et al. 2019), and the protein modification ontology PSI-MOD (Montecchi-Palazzi et al., 2008). We have contributed many new classes, corrections, and other revisions to all of these ontologies. For example, since 2018 PomBase curators have raised over 500 issues in the GO Consortium (GOC)’s GitHub tracker for ontology structure and content (https://github.com/geneontology/go-ontology/issues) and over 80 issues on the Mondo tracker (https://github.com/monarch-initiative/mondo/issues). We have also pioneered annotation quality control (QC) methods that are being adopted throughout the GOC. Notably, we developed a method to use observed co-annotation patterns to identify annotation outliers, and to build rules that allow automated outlier detection (Wood et al., 2020). Using this system, we have corrected thousands of annotations, and we collaborate with the GOC to continue rule development and to deploy a pipeline for error detection and reporting.

PomBase curators develop and maintain the Fission Yeast Phenotype Ontology (FYPO), a logically robust vocabulary that is designed for fission yeast but is also the leading candidate for further development into an ontology of cell-level phenotypes for all eukaryotes (Harris et al., 2013). FYPO currently comprises over 7500 terms, used in almost 97,000 annotations. FYPO development now uses the Ontology Development Kit (ODK; Matentzoglu 2021) for releases, which provides a release pipeline that seamlessly incorporates ontology reasoning, continuous integration (CI) checks, and generation of production files in OWL and OBO formats.

Our broad and deep ontology-based curation standardises data from large and small-scale publications, making a wide range of published data compatible with FAIR (Findable, Accessible, Interoperable and Reusable; Wilkinson et al. 2016) data sharing principles. Furthermore, our field-leading community curation project actively engages researchers in building the PomBase collection of FAIR-shared biological knowledge. We have recently described the insights gained from the project, including our experience with approaches that maximise participation, and the unanticipated added value that arises from co-curation by publication authors and professional curators (Lock et al., 2020). To date, we have assigned over 1800 publications to authors for curation, and have received over 975 submissions in response (54% response rate). Our online curation tool, Canto (Rutherford et al., 2014), has also been deployed for several other communities, including PHI-Base (Urban et al., 2020), FlyBase (Larkin et al., 2021), and the new *Schizosaccharomyces japonicus* MOD JaponicusDB (see Rutherford *et al*. in this issue).

Alongside its established, ongoing data stewardship activities, PomBase has introduced new features that enable biologists to integrate diverse molecular data into human-friendly summaries of biology. Below, we provide an overview of the new and updated features and describe how biologists can use new and existing data and tools to place their results into broader contexts. Taken together, PomBase activities give rise to a long-term collaboration with users to curate the knowledge gained from fission yeast research into an integrated overview of conserved cell biology to ensure that the resulting knowledge can be used to its full potential.

## Querying PomBase

PomBase provides robust, intuitive search interfaces to enable biologists to easily retrieve and combine data of diverse types from multiple sources.

### Simple search

A simple search, available in the header of every PomBase page, offers quick access to several commonly used PomBase pages. The search finds gene pages matching fission yeast names, synonyms, systematic IDs, UniProtKB accessions (The UniProt Consortium, 2020), or product descriptions. Human gene symbols (Tweedie et al., 2021) or *S. cerevisiae* gene names, ORF names, or IDs (Cherry et al., 2012) can also be used to find curated orthologs. Publication pages and ontology term pages can be retrieved using relevant IDs (PubMed or ontology IDs). Text searches generate autocomplete suggestions from matching gene product descriptions, ontology term names, and publication titles, author names, and dates.

### Advanced search

Since the reimplementation of the website in 2017, PomBase has provided an advanced search facility (https://www.pombase.org/query) that supports querying for gene sets based on a wide variety of criteria, including ontology annotations, gene product attributes (e.g. protein length, mass, domains or modifications), genomic location, conservation, etc. Complex queries can be constructed by combining single queries in the query history. The history, with links to result sets, is stored locally in the user’s web browser, and out-of-date results are highlighted and refreshed upon following result links. We have previously described links between ontology terms in query results, ontology term pages, and gene lists, which support data integration throughout the PomBase website (Lock et al., 2018).

Recent enhancements include improved phenotype querying, enhanced options for display and download of search results, and new tools for saving and sharing queries.

The search now provides an expanded set of query parameter options for phenotypes: experimental conditions can be used as search criteria for phenotype annotations, using the same condition descriptors as shown on PomBase web pages and in Canto. For example, a query can retrieve genes that show abnormal chromosome segregation mutant phenotypes specifically at high or low temperatures. Genes can also be selected based on the phenotypes of haploid or diploid, or single- or multi-locus, genotypes. For single-locus haploids, the expression level can be specified.

The display of search results is now highly customizable. Each query in the history has a link to a page of results that shows the count, query details, and a list of matching genes. By default, systematic IDs, names, and product descriptions are shown. The display can be customised by choosing additional columns (including a selection of gene expression data), and the list can be sorted on any column.

Similarly, selected data can be downloaded for genes in the list, via a popup that offers three sets of options:

- A “Tab delimited” option offers the same data as the results web page, for inclusion in a downloaded text file;
- A “Sequence” tab retrieves amino acid or nucleotide sequences in FASTA format, with checkboxes to select which items are included in the headers. When “Nucleotide” is selected, flanking sequence options similar to those on the gene page are available;
- A “GO annotation” tab downloads a file in GAF2.1 format (http://geneontology.org/docs/go-annotation-file-gaf-format-2.1/) that includes direct annotations (but not those inferred by transitivity) for the selected branch(es) of GO.

To facilitate query reuse and sharing, entries in the PomBase advanced search query history now show brief, user-editable query descriptions, and a toggle to show or hide additional details. All result pages from the advanced search now have a unique permanent URL that can be bookmarked and shared with colleagues. A set of “Commonly used queries” uses this system to provide convenient shortcuts to frequently sought data, such as all genes with disease associations (see below) or all protein-coding genes of unknown biological role.

### ID mapper

On a related note, we have developed an identifier mapper that retrieves *S. pombe* genes for a selection of different input ID types. Users can now find *S. pombe* genes using UniProtKB accessions, and retrieve manually curated orthologs for *S. cerevisiae* using standard gene names or ORF names, and for human using standard gene names or HUGO Gene Nomenclature Committee (HGNC; Tweedie et al. 2021) identifiers. *S. pombe* genes found via the ID mapper can be sent to the advanced search with a single click.

## From data to biological narratives

Much of PomBase’s recent work helps further our goal of enabling researchers to take data found in PomBase — from their own results combined with those of others — and build narratives that elucidate a biological topic with broad applicability.

### Intuitive annotation displays

In the PomBase reimplementation, we introduced display filtering on gene, publication, and ontology term pages for phenotype annotation tables, to narrow the list by broad phenotypic category, and (in the detailed view) by evidence. We have now added filtering for more annotation types, and expanded the set of filtering options. Display filtering is now available for all three branches of GO, and for qualitative or quantitative gene expression annotations. For GO and gene expression annotations, detailed views now offer filters for observations made during specific cell cycle phases, which use annotation extensions (Huntley et al., 2014) specifying the relevant phase. All of the above annotation types, as well as genetic and physical interactions, can also be filtered to distinguish between high-versus low-throughput experiments.

The display of protein features on gene pages has been updated to improve legibility, and provides an interactive graphical view in which mousing over any feature shows additional details in a tooltip and highlights the full entry in the accompanying table.

For ease of use, we have improved the appearance of PomBase pages on small screens such as tablets and smartphones.

### Comprehensive knowledge representation

To ensure that coverage of *S. pombe* research is current and complete, we periodically review our coverage of selected biological processes. We have recently performed comprehensive literature reviews and updated annotations in three broad areas of biology that are not intensively studied in fission yeast: tRNA metabolism, mitochondrial biology, and transmembrane transport. We have also reviewed a fourth topic, chromatin silencing, to bring annotations into alignment with substantial changes in how GO represents chromatin-mediated transcriptional regulation.

The *S. pombe* GO annotation corpus also now includes a set of annotations from the GOC’s Phylogenetic Annotation and INference Tool (PAINT) pipeline (Gaudet et al., 2011), which propagates GO annotation across all species based on protein family membership. Before incorporating PAINT annotations into PomBase, we reviewed the predictions made for *S. pombe* and reported errors to the GOC for over 500 protein families, improving the accuracy of the PAINT annotation corpus for all species. Like all GO annotations imported into PomBase from external sources, PAINT annotations are filtered for redundancy with experimentally supported annotations (Lock et al., 2018).

Qualitative gene expression annotations now use an expanded set of descriptors, accommodating ribosomal density data as well as more precisely defined RNA and protein level terms.

To ensure coverage of high-throughput experiments, we have continued to build the collection of data tracks, and associated well-curated metadata, in the PomBase JBrowse (Buels et al., 2016) instance; the browser now hosts 330 tracks from 28 publications. To improve browser data visibility, gene pages now display JBrowse images that have clickable features and a link to open the fully functional browser in a new tab or window. Data tracks from datasets hosted in the PomBase genome browser can also now be browsed, selected, and loaded from their respective publication pages.

PomBase is now a Partner Database with microPublication, whose remit is to publish “brief, novel findings, negative and/or reproduced results, and results which may lack a broader scientific narrative” (https://www.micropublication.org/; Raciti et al. 2018). This partnership thus enables fission yeast researchers to publish individual experimental results that have not been included in traditional publications, thereby building a collection of *S. pombe* data that fill gaps in available datasets.

Publication pages are now also available for reviews and methods papers as well as microPublications.

### Ontology Slims

Ontology subsets can provide overviews of annotation across a gene set or the whole genome. PomBase has long provided the fission yeast Biological Process (BP) GO slim, a subset of the GO BP ontology designed to cover as many annotated gene products as possible, while remaining informative about the gene product’s physiological role in the cell (Lock et al., 2018; Wood et al., 2019). To complement the BP slim, we have now created three new ontology subsets. The fission yeast Molecular Function (MF) and Cellular Component (CC) slims summarise the MF and CC branches of GO respectively; together, all three GO slims provide a simple yet comprehensive summary of *S. pombe*’s biological capabilities by grouping gene products using broad classifiers from the full breadth of GO.

The third new ontology subset is drawn from Mondo, and gives an overview of genes with human orthologs implicated in disease. To accompany the Mondo slim, we have improved coverage for gene curation that associates disease descriptors with fission yeast orthologs of human disease-causing genes; over 1400 *S. pombe* genes now have curated disease associations. PomBase curators also collaborate with Mondo to improve its disease classification, especially in areas relevant to fission yeast disease gene associations.

All of the PomBase ontology slims are integrated into annotation displays and querying. For each slim, a summary page lists the terms and IDs with links to ontology term pages, and provides a genome-wide overview of annotated genes. In addition to the number of genes annotated to each slim term, the slim page identifies sets of genes that are not annotated to any term in the ontology, or are annotated only to terms that do not have paths in the ontology to any slim term. On gene pages, annotation tables show slim terms applicable to a gene in the headers. On all pages where they appear, gene lists are linked to the advanced search, allowing the gene list to be combined with any other results in a new query. Slim annotations can also be retrieved for any advanced search result list.

We also maintain an up-to-date list of protein-coding *S. pombe* genes that are broadly conserved in eukaryotes (present in vertebrates), but have not been assigned an informative role from the GO BP slim (https://www.pombase.org/status/priority-unstudied-genes; Wood et al. 2019).

### QuiLT

The Quick Little Tool (QuiLT) is a new feature that allows users to view multiple types of annotation in a single graphical display. Inspired by our analysis of conserved unstudied proteins (Wood et al., 2019), QuiLT generates a graphic for any gene list uploaded or obtained in advanced search results. QuiLT visualisation is also linked to PomBase pages that list genes annotated to an ontology term, and on the “Priority unstudied genes” page (see above). QuiLT has display options for deletion viability, presence or absence of budding yeast orthologs, presence or absence of human orthologs, annotation from each branch of GO, characterisation status for protein-coding genes, taxonomic distribution, and protein length. The display is interactive, allowing users to highlight subsets of the list, filter the display, toggle annotation types on and off, reorder the list to focus on features of most interest, and download the image.

Figure 1 shows QuiLT visualisation of genes that are conserved in vertebrates, and were found to be associated with chronological lifespan in a recent study (Romila et al., 2021). The authors compared their list of long-lived mutants with all genes previously associated with the phenotype ‘increased viability in stationary phase’ (FYPO:0001309) to uncover novel ageing-associated genes; among the latter genes, they then readily identified conserved genes using the catalog of conserved unstudied genes described above, as well as genes associated with diseases in human.

**Fig. 1.**
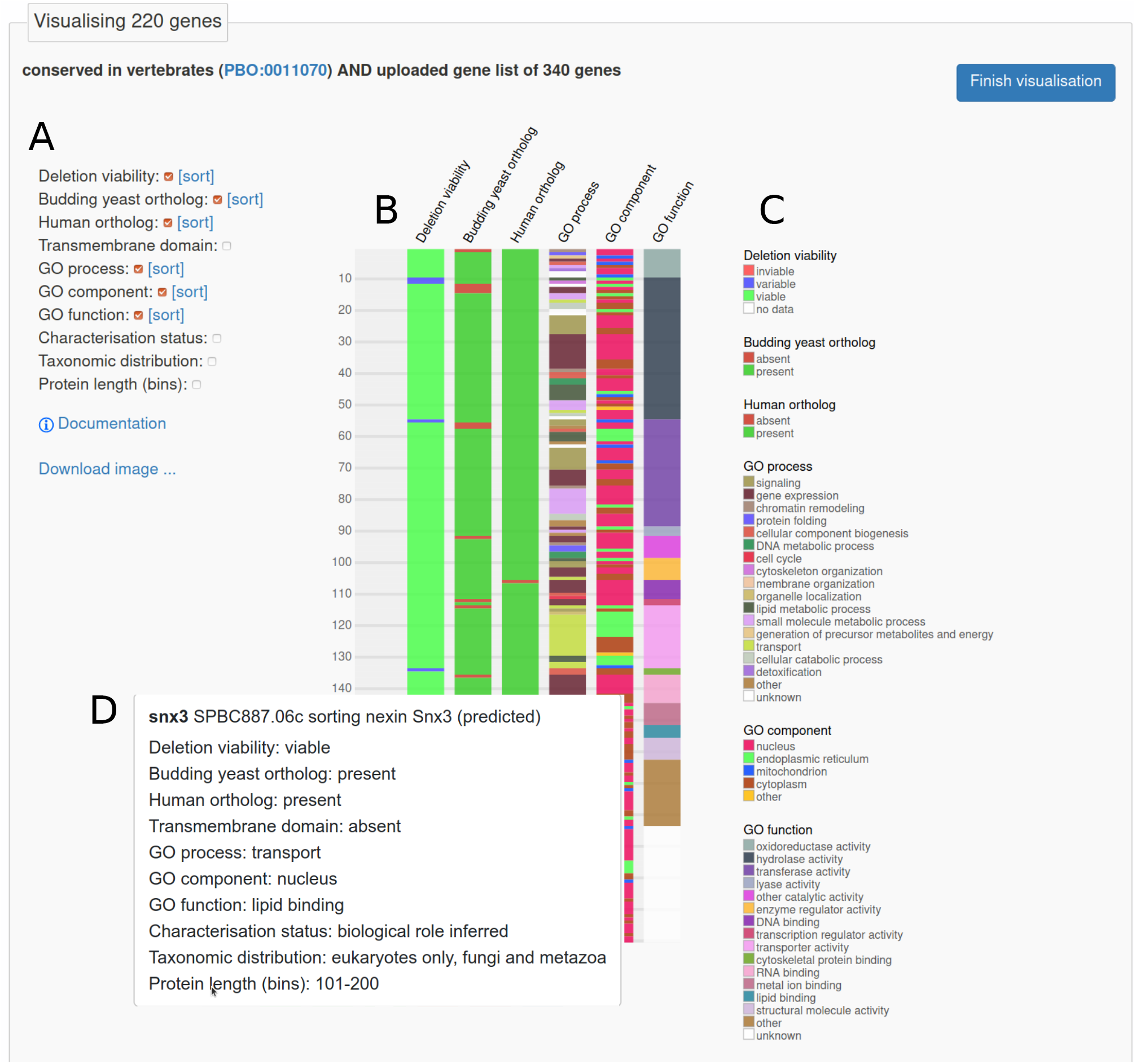
Quick Little Tool (QuiLT) visualisation of 220 *S. pombe* genes showing viability of the deletion mutant, presence or absence of orthologs in human and budding yeast, and GO annotation. Romila et al. (2021) identified 340 genes for which deletion increased chronological lifespan; the visualisation uses 220 of those genes annotated in PomBase as conserved in vertebrates. (A) Interactive display controls show or hide annotation types, and sort on any visible column. The image can be downloaded in SVG format. (B) Four annotation types (transmembrane domain, characterisation status, taxonomic distribution, and protein length) were hidden, and the display was sorted on GO molecular function, yielding the graphic shown. (C) Key to colours for each visible annotation type in (B). (D) Mousing over blocks in the graphic (B) displays additional details in pop-ups. The leftmost column represents one row per gene, and the pop-up shows details including both visible and hidden annotation values. The interactive version of the visualisation is available online at https://www.pombase.org/vis/from/id/d737355c-00b1-4692-bb13-6f9752b28436.

### Pathways

A new gene page section, “Molecular pathway”, connects genes in PomBase to depictions of biochemical and signalling pathways. At present this section is shown for any gene that appears in a pathway entry in the Kyoto Encyclopedia of Genes and Genomes (KEGG) database (Kanehisa and Goto, 2000; Kanehisa, 2019; Kanehisa et al., 2021), linking to the relevant KEGG page(s) and to a PomBase page listing all genes connected to the pathway. The Molecular pathway section will figure prominently in future development of data integration in PomBase (see below).

## Future Directions

We will continue all of the well-established activities described above, ensuring that PomBase presents comprehensive, quality-controlled, curated knowledge from large- and small-scale *S. pombe* experiments. In addition, a diverse set of enhancements are under development.

To improve the accuracy and coverage of sequence feature annotation, a comprehensive overhaul of 5’ and 3’ untranslated region (UTR) lengths is in progress, and we will gradually add observed transcript isoforms to more genes. The whole genome sequence will also be updated to fill gaps, correct errors, and incorporate new telomeric sequence data, and resubmitted to the International Nucleotide Sequence Database Collaboration (INSDC; https://www.insdc.org/) databases.

We will continue to enhance PomBase’s simple and advanced search tools with new features and capabilities, such as better handling of genes with multiple transcript isoforms, improved querying for phenotype conditions, and import/export of query histories. We also plan to investigate options, such as user accounts, to enable researchers to configure PomBase page displays, browser settings, etc. and to make saved settings and query histories more portable.

Most importantly, we will focus effort on building richer, more comprehensive connections among curated data of all types. Notably, we will adapt *S. pombe* GO annotations to follow the “GO Causal Activity Modeling” principles established by the GO Consortium for GO-CAMs (Thomas et al., 2019), using our large, detailed body of existing GO annotations. Manually curated *S. pombe* GO annotations already include a rich set of extensions that capture reaction substrates, effector–target connections, high-confidence physical interactions, complexes, and regulatory effects. Converting these extended annotations to GO-CAM models will improve PomBase’s representation of how protein activities are connected into pathways and how pathways are connected to each other. Pathway diagrams generated by software using GO-CAM data will be shown in the Molecular pathway section of gene pages. Furthermore, by integrating GO-CAM-based pathways with curated information about which genes and pathways relate to human diseases, and what proteins remain unstudied across species, PomBase will enable users to develop an emergent understanding of cell biology relevant throughout the eukaryotes. Biological stories crafted from fission yeast data will thus shed light on all biological systems.

## Data Availability

PomBase has used open-source tools and code to develop a modular, customizable and readily reusable system that supports daily data updates and an intuitive web-based user interface (Lock et al., 2018). All PomBase code is available from the PomBase GitHub organisation (https://github.com/pombase), where each major aspect of the project — including curation, website, Chado database, FYPO, and Canto — has a dedicated repository.

In addition to the web page displays and query result downloads described above, PomBase data can be downloaded in bulk from the website (see https://www.pombase.org/datasets and https://www.pombase.org/data). We have recently switched all downloads to use the HTTPS protocol, superseding FTP. Available downloads include nightly dumps and monthly snapshots of the entire Chado curation database, and a range of specific curated datasets, including GO annotations, single-allele phenotypes, protein modifications, high-confidence physical interactions, and manually curated ortholog lists.

## Acknowledgements

PomBase thanks the fission yeast community for contributing datasets and literature curation, providing valuable feedback on the website, and for their continued support. We thank Nico Matentzoglu for assistance with deploying ODK to manage FYPO development, and, along with David Osumi-Sutherland and other uPheno project contributors, for advice on logical definition patterns for FYPO terms. Daniela Raciti and the rest of the microPublications team have provided essential assistance setting up *S. pombe* microPublications, and offer ongoing advice and support. We are grateful to Pascale Gaudet for acting on our recommendations for improving GO annotations and ontology content. We also thank Jon Hollis and others at the Babraham Institute for hosting and support of the PomBase website.

## References

R. Buels, E. Yao, C. M. Diesh, R. D. Hayes, M. Munoz-Torres, G. Helt, D. M. Goodstein, C. G. Elsik, S. E. Lewis, L. Stein, and I. H. Holmes. JBrowse: a dynamic web platform for genome visualization and analysis. Genome Biol, 17:66, 2016.

J. M. Cherry, E. L. Hong, C. Amundsen, R. Balakrishnan, G. Binkley, E. T. Chan, K. R. Christie, M. C. Costanzo, S. S. Dwight, S. R. Engel, D. G. Fisk, J. E. Hirschman, B. C. Hitz, K. Karra, C. J. Krieger, S. R. Miyasato, R. S. Nash, J. Park, M. S. Skrzypek, M. Simison, S. Weng, and E. D. Wong. Saccharomyces Genome Database: the genomics resource of budding yeast. Nucleic Acids Res, 40(Database issue):D700–D705, 2012.

K. Eilbeck, S. E. Lewis, C. J. Mungall, M. Yandell, L. Stein, R. Durbin, and M. Ashburner. The Sequence Ontology: a tool for the unification of genome annotations. Genome Biol, 6 (5):R44, 2005.

P. Gaudet, M. S. Livstone, S. E. Lewis, and P. D. Thomas. Phylogenetic-based propagation of functional annotations within the Gene Ontology consortium. Brief Bioinform, 12 (5):449–462, 2011.

M. A. Harris, A. Lock, J. Bähler, S. G. Oliver, and V. Wood. FYPO: the fission yeast phenotype ontology. Bioinformatics, 29(13):1671–1678, 2013.

R. P. Huntley, M. A. Harris, Y. Alam-Faruque, J. A. Blake, S. Carbon, H. Dietze, E. C. Dimmer, R. E. Foulger, D. P. Hill, V. K. Khodiyar, A. Lock, J. Lomax, R. C. Lovering, P. Mutowo-Meullenet, T. Sawford, K. V. Auken, V. Wood, and C. J. Mungall. A method for increasing expressivity of Gene Ontology annotations using a compositional approach. BMC Bioinformatics, 15:155, 2014.

M. Kanehisa. Toward understanding the origin and evolution of cellular organisms. Protein Sci, 28(11):1947–1951, 2019.

M. Kanehisa and S. Goto. KEGG: kyoto encyclopedia of genes and genomes. Nucleic Acids Res, 28(1):27–30, 2000.

M. Kanehisa, M. Furumichi, Y. Sato, M. Ishiguro-Watanabe, and M. Tanabe. KEGG: integrating viruses and cellular organisms. Nucleic Acids Res, 49(D1):D545–D551, 2021.

A. Larkin, S. J. Marygold, G. Antonazzo, H. Attrill, G. dos Santos, P. V. Garapati, J. L. Goodman, L. S. Gramates, G. Millburn, V. B. Strelets, C. J. Tabone, J. Thurmond, and the FlyBase Consortium. FlyBase: updates to the Drosophila melanogaster knowledge base. Nucleic Acids Res, 49(Database issue):D899–D907, 2021.

A. Lock, K. Rutherford, M. A. Harris, J. Hayles, S. G. Oliver, J. Bähler, and V. Wood. PomBase 2018: user-driven reimplementation of the fission yeast database provides rapid and intuitive access to diverse, interconnected information. Nucleic Acids Research, 47(D1):D821–D827, 2018. doi: 10.1093/nar/gky961. URL http://dx.doi.org/10.1093/nar/gky961.

A. Lock, M. A. Harris, K. Rutherford, J. Hayles, and V. Wood. Community curation in PomBase: enabling fission yeast experts to provide detailed, standardized, sharable annotation from research publications. Database (Oxford), 2020:baaa028, 2020. ISSN 1758-0463. doi: 10.1093/database/baaa028. URL https://doi.org/10.1093/database/baaa028.

N. Matentzoglu. INCATools/ontology-development-kit: June 2020 release, June 2021. URL https://doi.org/10.5281/zenodo.4973944.

L. Montecchi-Palazzi, R. Beavis, P.-A. Binz, R. J. Chalkley, J. Cottrell, D. Creasy, J. Shofstahl, S. L. Seymour, and J. S. Garavelli. The PSI-MOD community standard for representation of protein modification data. Nat Biotechnol, 26(8):864–866, 2008.

C. J. Mungall, J. A. McMurry, S. Köhler, J. P. Balhoff, C. Borromeo, M. Brush, S. Carbon, T. Conlin, N. Dunn, M. Engelstad, E. Foster, J. P. Gourdine, J. O. B. Jacobsen, D. Keith, B. Laraway, S. E. Lewis, J. NguyenXuan, K. Shefchek, N. Vasilevsky, Z. Yuan, N. Washington, H. Hochheiser, T. Groza, D. Smedley, P. N. Robinson, and M. A. Haendel. The Monarch Initiative: an integrative data and analytic platform connecting phenotypes to genotypes across species. Nucleic Acids Res, 45(D1):D712–D722, 2017.

D. A. Natale, C. N. Arighi, J. A. Blake, J. Bona, C. Chen, S.-C. Chen, K. R. Christie, J. Cowart, P. D’Eustachio, A. D. Diehl, H. J. Drabkin, W. D. Duncan, H. Huang, J. Ren, K. Ross, A. Ruttenberg, V. Shamovsky, B. Smith, Q. Wang, J. Zhang, A. El-Sayed, and C. H. Wu. Protein Ontology (PRO): enhancing and scaling up the representation of protein entities. Nucleic Acids Res, 45(D1):D339–D346, 2017.

R. Oughtred, J. Rust, C. Chang, B.-J. Breitkreutz, C. Stark, A. Willems, L. Boucher, G. Leung, N. Kolas, F. Zhang, S. Dolma, J. Coulombe-Huntington, A. Chatr-Aryamontri, K. Dolinski, and M. Tyers. The BioGRID database: A comprehensive biomedical resource of curated protein, genetic, and chemical interactions. Protein Sci, 30(1): 187–200, 2021.

D. Raciti, K. Yook, T. W. Harris, T. Schedl, and P. W. Sternberg. Micropublication: incentivizing community curation and placing unpublished data into the public domain. Database (Oxford), 2018:bay013, 2018.

C. A. Romila, S. Townsend, M. Malecki, S. Kamrad, M. Rodríguez-López, O. Hillson, C. Cotobal, M. Ralser, and J. Bähler. Barcode sequencing and a high-throughput assay for chronological lifespan uncover ageing-associated genes in fission yeast. Microb Cell, 8(7):146–160, 2021.

K. M. Rutherford, M. A. Harris, A. Lock, S. G. Oliver, and V. Wood. Canto: an online tool for community literature curation. Bioinformatics, 30(12):1791–1792, 2014.

K. A. Shefchek, N. L. Harris, M. Gargano, N. Matentzoglu, D. Unni, M. Brush, D. Keith, T. Conlin, N. Vasilevsky, X. A. Zhang, J. P. Balhoff, L. Babb, S. M. Bello, H. Blau, Y. Bradford, S. Carbon, L. Carmody, L. E. Chan, V. Cipriani, A. Cuzick, M. Della Rocca, N. Dunn, S. Essaid, P. Fey, C. Grove, J.-P. Gourdine, A. Hamosh, M. Harris, I. Helbig, M. Hoatlin, M. Joachimiak, S. Jupp, K. B. Lett, S. E. Lewis, C. McNamara, Z. M. Pendlington, C. Pilgrim, T. Putman, V. Ravanmehr, J. Reese, E. Riggs, S. Robb, P. Roncaglia, J. Seager, E. Segerdell, M. Similuk, A. L. Storm, C. Thaxon, A. Thessen, J. O. B. Jacobsen, J. A. McMurry, T. Groza, S. Köhler, D. Smedley, P. N. Robinson, C. J. Mungall, M. A. Haendel, M. C. Muñoz-Torres, and D. Osumi-Sutherland. The Monarch Initiative in 2019: an integrative data and analytic platform connecting phenotypes to genotypes across species. Nucleic Acids Research, 48(D1):D704–D715, 11 2019. ISSN 0305-1048. doi: 10.1093/nar/gkz997. URL https://doi.org/10.1093/nar/gkz997.

The Gene Ontology Consortium. Gene Ontology: tool for the unification of biology. Nat Genet, 25(1):25–29, 2000.

The Gene Ontology Consortium. The Gene Ontology resource: enriching a GOld mine. Nucleic Acids Res, 49(D1):D325–D334, 2021.

The UniProt Consortium. UniProt: the universal protein knowledgebase in 2021. Nucleic Acids Research, 49(D1): D480–D489, 2020. ISSN 0305-1048. doi: 10.1093/nar/gkaa1100.

P. D. Thomas, D. P. Hill, H. Mi, D. Osumi-Sutherland, K. V. Auken, S. Carbon, J. P. Balhoff, L.-P. Albou, B. Good, P. Gaudet, S. E. Lewis, and C. J. Mungall. Gene Ontology Causal Activity Modeling (GO-CAM) moves beyond GO annotations to structured descriptions of biological functions and systems. Nat Genet, 51(10):1429–1433, 2019.

S. Tweedie, B. Braschi, K. Gray, T. E. M. Jones, R. L. Seal, B. Yates, and E. A. Bruford. Genenames.org: the HGNC and VGNC resources in 2021. Nucleic Acids Res, 49(D1): D939–D946, 2021.

M. Urban, A. Cuzick, J. Seager, V. Wood, K. Rutherford, S. Y. Venkatesh, N. D. Silva, M. C. Martinez, H. Pedro, A. D. Yates, K. Hassani-Pak, and K. E. Hammond-Kosack. PHI-base: the pathogen-host interactions database. Nucleic Acids Res, 48(D1):D613–D620, 2020.

M. D. Wilkinson, M. Dumontier, I. J. Aalbersberg, G. Appleton, M. Axton, A. Baak, N. Blomberg, J.-W. Boiten, L. B. da Silva Santos, P. E. Bourne, J. Bouwman, A. J. Brookes, M. Clark, T. Crosas, O. Dillo, I. Dumon, S. Edmunds, C. R. Evelo, T. Finkers, A. Gonzalez-Beltran, A. J. G. Gray, P. Groth, C. Goble, J. S. Grethe, J. Heringa, P. A. C. ‘t Hoen, T. Hooft, R. Kuhn, R. Kok, J. Kok, S. J. Lusher, M. E. Martone, A. Mons, A. L. Packer, B. Persson, M. Rocca-Serra, P. Roos, R. van Schaik, S.-A. Sansone, E. Schultes, T. Sengstag, T. Slater, G. Strawn, M. A. Swertz, M. Thompson, J. van der Lei, E. van Mulligen, J. Velterop, A. Waagmeester, P. Wittenburg, K. Wolstencroft, J. Zhao, and B. Mons. The FAIR Guiding Principles for scientific data management and stewardship. Sci Data, 3:160018, 2016.

V. Wood, A. Lock, M. A. Harris, K. Rutherford, J. Bähler, and S. G. Oliver. Hidden in plain sight: what remains to be discovered in the eukaryotic proteome? Open Biol, 9(2): 180241, 2019.

V. Wood, S. Carbon, M. A. Harris, A. Lock, S. R. Engel, D. P. Hill, K. Van Auken, H. Attrill, M. Feuermann, P. Gaudet, R. C. Lovering, S. Poux, K. M. Rutherford, and C. J. Mungall. Term Matrix: a novel Gene Ontology annotation quality control system based on ontology term co-annotation patterns. Open Biol, 10:200149, 2020.

